# Modelling the impact of climate change on the distribution and abundance of tsetse in Northern Zimbabwe

**DOI:** 10.1101/2020.07.03.186338

**Authors:** Joshua Longbottom, Cyril Caminade, Harry S. Gibson, Daniel J. Weiss, Steve Torr, Jennifer S. Lord

**Author notes:** Corresponding author: Joshua Longbottom, Department of Vector Biology, Liverpool School of Tropical Medicine, Liverpool, L3 5QA, UK.

## Abstract

**Background:** Climate change is predicted to impact the transmission dynamics of vector-borne diseases. Tsetse flies (*Glossina*) transmit species of *Trypanosoma* that cause human and animal African trypanosomiasis. A previous modelling study showed that temperature increases between 1990 and 2017 can explain the observed decline in abundance of tsetse at a single site in the Mana Pools National Park of Zimbabwe. Here, we apply a mechanistic model of tsetse population dynamics to predict how increases in temperature may have changed the distribution and relative abundance of *Glossina pallidipes* across northern Zimbabwe.

**Methods:** Local weather station temperature measurements were previously used to fit the mechanistic model to longitudinal *G. pallidipes* catch data. To extend the use of the model, we converted MODIS land surface temperature to air temperature, compared the converted temperatures with available weather station data to confirm they aligned, and then re-fitted the mechanistic model using *G. pallidipes* catch data and air temperature estimates. We projected this fitted model across northern Zimbabwe, using simulations at a 1 km × 1 km spatial resolution, between 2000 to 2016.

**Results:** We produce estimates of relative changes in *G. pallidipes* mortality, larviposition, emergence rates and abundance, for northern Zimbabwe. Our model predicts decreasing tsetse populations within low elevation areas in response to increasing temperature trends during 2000-2016. Conversely, we show that high elevation areas (>1000 M.A.S.L), previously considered too cold to sustain tsetse, may now be climatically suitable.

**Conclusions:** The results of this research represent the first regional-scale assessment of temperature related tsetse population dynamics, and the first high spatial-resolution estimates of this metric for northern Zimbabwe. Our results suggest that tsetse abundance may have declined across much of the Zambezi valley in response to changing climatic conditions during the study period. Future research including empirical studies is planned to improve model accuracy and validate predictions for other field sites in Zimbabwe.

## Background

Human African trypanosomiasis (HAT, also referred to as ‘sleeping sickness’), is a neglected tropical disease caused by subspecies of *Trypanosoma brucei*. The disease exists as two differing pathologies: Gambian sleeping sickness (g-HAT), caused by *Trypanosoma brucei gambiense*, is generally considered to be an anthroponosis and, with no vaccines or prophylactic drugs existing, disease control efforts rely primarily on active/passive case detection and treatment of human cases, combined sometimes with vector control [1]. Rhodesian sleeping sickness (r-HAT), caused by *T. b. rhodesiense* [2], is a zoonosis with wild animals and cattle acting as reservoir hosts. While therapeutic drugs for r-HAT exist, mass screening and treatment of humans has little effect on transmission between reservoir hosts. Accordingly, control of r-HAT relies on vector control and treatment of domestic reservoir hosts with trypanocides [3]. Despite a lower contribution to the overall number of HAT cases (3% [4]), r-HAT has a more complicated pathology, causing acute infection ultimately resulting in death.

Recent discussions have resulted in the inclusion of r-HAT in the 2030 neglected tropical disease roadmap, with a target of no endemic areas reporting > 1 HAT case per 10 000 people per year (average of 5 years) by 2030 [5]. Both forms of HAT are limited by the spatial distribution of the tsetse fly vector, and as such HAT is endemic within 36 countries in sub-Saharan Africa [2, 6]. Additionally, alongside human burden, tsetse transmit species of Trypanosoma which cause animal African trypanosomiasis (AAT) which kills about 1 million cattle per year, posing a further risk to upwards of 55 million cattle [7, 8].

A thorough understanding of the ecology of tsetse is essential for implementing effective control measures [9, 10]. Tsetse population dynamics vary spatially, and environmental drivers such as temperature influence key aspects of tsetse ecology and demography, including vector survival, development and fecundity, and ultimately spatial distribution and density [11, 12]. The effect of seasonal variations in temperature on adult and pupal survival has been widely studied within the laboratory and field [13, 14, 15, 16]. Any factors which result in a change of tsetse vital rates, particularly factors ultimately altering population age structure and abundance, can in turn, affect disease risk. As abiotic conditions fluctuate in both space and time, particularly in the face of global climate change, there is a need to understand how spatio-temporal environmental variation drives *Glossina* population dynamics; high spatial resolution predictive mapping of population dynamics could elucidate this problem and enable enhanced tsetse surveillance and control.

Climate change has complex implications for both vector and disease distributions. Global temperatures were 1.31°C greater in 2017 than the 20^th^ century average [17], with estimates from the Intergovernmental Panel on Climate Change suggesting temperature increases will likely be in the range of 0.3-0.7°C between 2016 and 2035, even under the most optimistic of scenarios [18]. Lord et al., used a -27-year time-series of tsetse abundance from the Mana Pools National Park, Zimbabwe, to show that temperature increases of around 2°C between 1975 and 2017 can explain a >90% decrease in tsetse abundance at that location [19]. Within Northern Zimbabwe, cases of reported r-HAT have also declined, with only five reported cases within the last three years (2015-2017, range 1-3 cases/year) compared with 13 cases during 2012-2014 (range, 1-9 cases/year) [20]. Whether the decline in incidence of r-HAT is related to changes in tsetse populations is unclear.

The effect of temperature on the distribution of tsetse throughout the rest of Northern Zimbabwe is currently unknown due to limited sampling/dissemination of sampling results. Catch data from Rekomitjie field station exists as one of the most comprehensive longitudinal datasets of tsetse count data available to date. Rekomitjie lies within the Mana Pools National Park and tsetse populations have not been subjected to any control measures or gross environmental change related to farming or human settlement for >60 years. The catch data obtained from Rekomitjie, therefore, are highly indicative of the response of tsetse populations to abiotic changes at the field site location, and therefore form a suitable dataset for the construction of a temperature-dependent population dynamic model (as shown by Lord et al. [19]).

This study aimed to expand on the approach used by Lord et al. [19], to model potential changes in *G. pallidipes* populations across Northern Zimbabwe. By spatially projecting a model of tsetse population dynamics, we aimed to identify locations where viable numbers of tsetse may persist, or where environments may have become more suitable with respect to temperature, allowing for targeted vector monitoring, control and improved predictions of r-HAT risk for this region.

## Methods

### Temperature data

Our analyses focussed on Northern Zimbabwe (bounding box: 24.99°E, 19.00°S, 34.00°E, 14.99°S) (Fig. 1). For this area, 1 × 1 km resolution land-surface temperature (LST) data from the Moderate Resolution Imaging Spectroradiometer (MODIS), gap-filled to remove cloud cover as described in Weiss *et al*. [21, 22], were obtained for each month between March 2000 and December 2016. The processed surfaces included separate measurements for mean monthly daytime LST (*LST*_*day*_) and mean monthly night-time LST (*LST*_*night*_) for each grid cell. Each surface was derived from multiple eight-day composites, with each cell containing an average value generated from between two to eight measurements depending on data quality [23]. Daytime measurements represent temperatures at c. 10:30AM local time and night-time measurements represent temperatures at c. 10:30PM local time due to the overhead passing of the MODIS satellite. The two differing surfaces were combined to produce a mean monthly LST surface (*LST*_*mean*_) and a difference surface (*LST*Δ) that captured the diurnal temperature flux.

**Figure 1.**
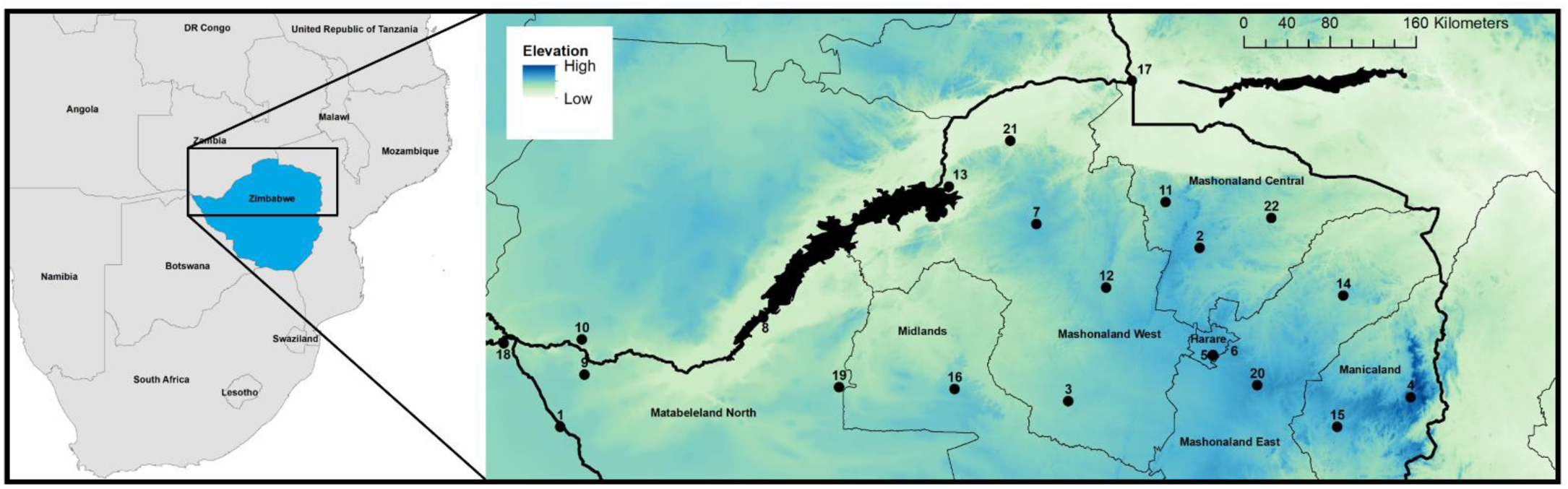
Study area. Numbered locations represent the network of weather stations used to assess accuracy of MODIS adjustment from land-surface temperature to air temperature. Elevation (m) is used as the background dataset, black polygons represent waterbodies and black lines represent administrative boundaries (thin lines: provincial boundary, thick lines: international borders).

We obtained daily minimum, mean and maximum temperatures from the weather station at Rekomitjie Research Station, Zimbabwe, from October 1959 to July 2017. For the period over which MODIS data were available, we derived monthly mean temperatures. We then calculated the difference between monthly MODIS LST and air temperatures from the weather station for each month between March 2000 and December 2016.

To align the weather station and MODIS temperatures, we converted *LST*_*day*_ and *LST*_*night*_ to minimum and maximum air temperatures as per Weiss *et al*. [22]. Estimates of day length, in hours, for each cell within the study area were computed as described in Forsythe *et al*. [24]. Using these estimates and an index for the number of days per month, the mean day length (*DL*_*mean*_) per cell was calculated for each month. These monthly outputs, and the *LST*_*day*_, *LST*_*night*_ and *LST*Δ surfaces were incorporated with regression coefficients identified by Weiss *et al*. [22], with a corrected intercept (pers. comm.) to compute maximum (*Air*_*max*_) (EQ1) and minimum (*Air*_*min*_) (EQ2) air temperatures. The mean air temperature (*Air*_*mean*_) estimates were then calculated as the average of *Air*_*max*_ and *Air*_*min*_.

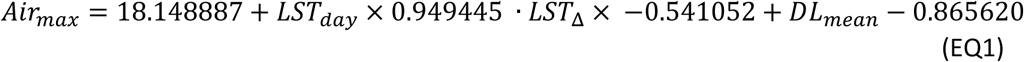

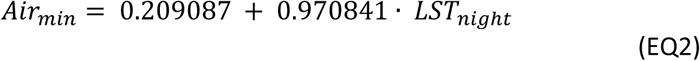

(EQ2)In order to validate the conversion of the MODIS LST data to air temperature, data were obtained from the Global Surface Summary of the Day [25] for 21 weather stations within the study extent for 2000-2016 (Fig. 1). Monthly mean air temperatures were calculated for each station which had two or more measurements per day, and 15 or more daily measurements per month. We compared the measured air temperature data from stations with the MODIS-derived air temperature (*Air*_*mean*_) using linear regression with *Air*_*mean*_ as the explanatory variable.

### Tsetse population dynamics

Within Zimbabwe, the primary r-HAT vectors are *G. morsitans morsitans* and *G. pallidipes. Glossina pallidipes* occupies a wide range throughout much of the Zambezi Valley, proving to be an effective bridge vector feeding on wild game, domestic cattle and humans [26, 27]. As fully described in Lord *et al*.[19], since 1966, daily collections of female *G. pallidipes* from stationary oxen have been performed at Rekomitjie field station (location 21, Fig. 1), to monitor insecticide efficacy [28]. Throughout the 1960s to the 1980s, a quota of 50 flies a day was set concurrent with the minimum expected catch at that time. Records on the reported number of flies caught per day exist from 1990 to present day. Due to the temporal availability of the MODIS data, we used catch data only from March 2000 to December 2016 for this study.

Tsetse are different from most biting flies in that they retain a single fertilised egg within their uterus where it develops into a third-instar larva in about nine days [16]. The larva is deposited by the female and burrows quickly into the ground, where it pupates and spends c. 30 days before emerging as an adult fly. The adult female produces its first offspring about 15 days after emergence, so that the minimum generation time is around 45 to 50 days, and thereafter the female produces just one larva every nine days [29]. This very slow rate of reproduction makes tsetse populations sensitive to relatively small increases in mortality, it being estimated that an added mortality of just 3% per day results in population extinction in a year or so if there is no immigration [16, 30, 31].

There is a wealth of publicly available data for the way that vital rates of tsetse respond to differing environmental conditions studied in both the field and the laboratory. Using such data for *G. pallidipes*, Lord et al. produced and collated equations relating daily temperature to daily female adult mortality rate (*μ*_*A*_) (Eq. 3), daily female pupal mortality rate (*μ*_*p*_) (Eq. 4), daily pupal development rate (*β*) (Eq. 5), and daily larviposition rate (*ρ*) (Eqs. 6 & 7). These temperature-dependent processes and a density-dependent mortality coefficient (*δ*), were used in a set of three ordinary differential equations (ODE) describing tsetse population dynamics [19] (Fig. 2).

**Figure 2.**
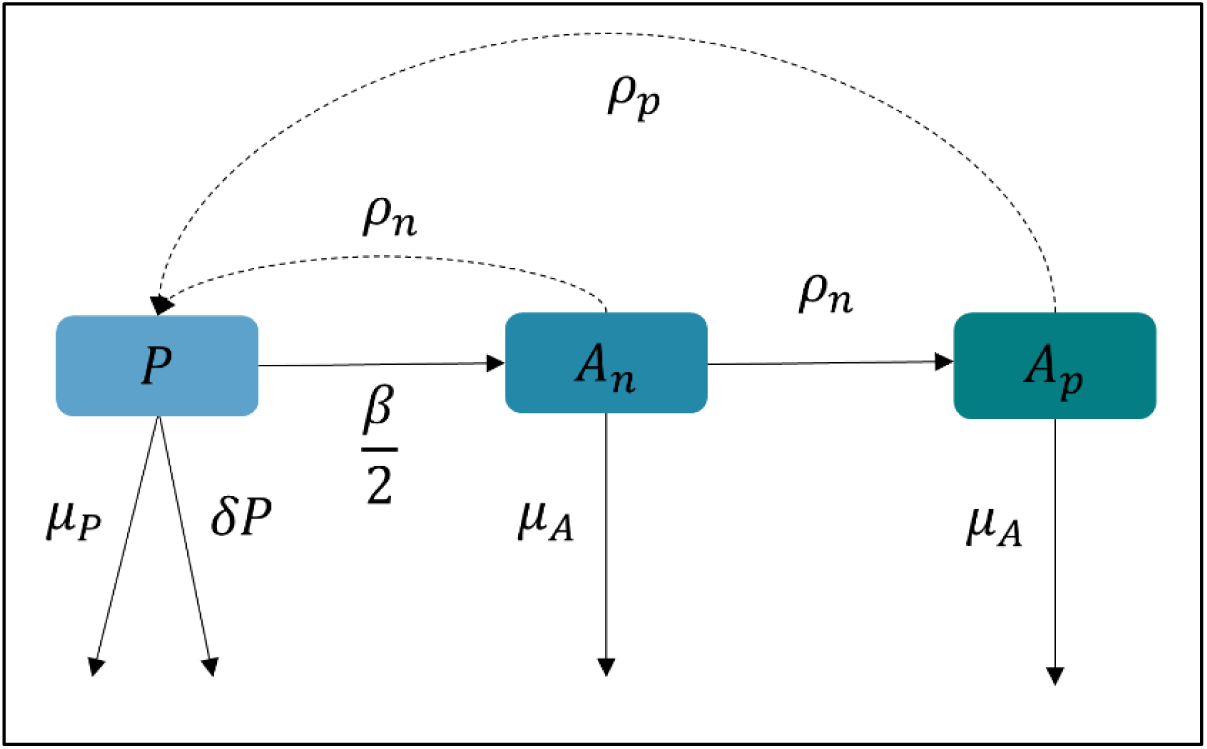
Pictorial representation of the ODE model described in Lord *et al*. [19]. The ODE model considers three states: *P*, pupae; *A*_*n*_, nulliparous adults, and *A*_*p*_, parous adults.

At each spatial location, pupae are produced by nulliparous (*A*_*n*_) and parous (*A*_*p*_) adult females at rates *ρ*_*n*_ and *ρ*_*p*_ respectively. As the number of adult females at a location is also temperature-dependent, the number of nulliparous and parous individuals at each location can be considered by the following: losses from the pupal stage are due to (i) pupae emerging as nulliparous adults (*A*_*n*_) at rate *β*; (ii) density-dependent mortality, with coefficient *δ* and (iii) pupal mortality *μ*_*P*_. Losses from the nulliparous adult stage are due to first larviposition, occurring at rate *ρ*_*n*_ and adult mortality *μ*_*A*_. For this study, the adult mortality rate is assumed to be equal for both nulliparous and parous females. The mentioned states form the structure of the model (Fig 2), fully described by Lord et al. [19].

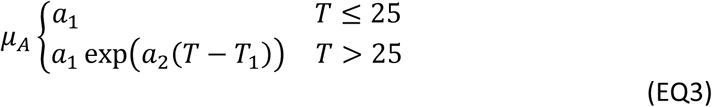

Where *T* is temperature in °C. *T*_1_is not a parameter but a constant set to 25 to ensure that *a*_2_ is within a convenient range.

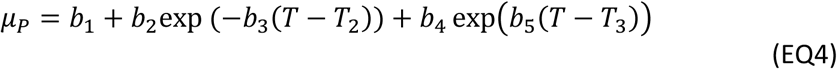

Where *T* is temperature in °C. *T*_2_ and *T*_3_ are not parameters but are constants chosen to ensure that the coefficients *b*_3_ and *b*_5_ are within a convenient range, and were set to 16°C and 32°C respectively.

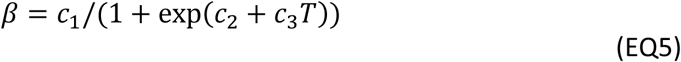

Where for females, the fitted estimates were *c*_1_ = 0.05884, *c*_2_ = 4.8829, and *c*_3_ = –0.2159.

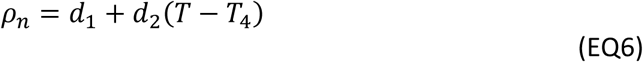

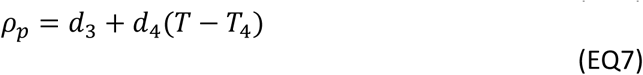

Where *ρ*_*p*_ represents the larviposition rate for parous adults and *ρ*_*n*_ represents the larviposition rate for nulliparous adults. *T*_4_ was set to 24°C, *d*_1_ = 0.061, *d*_2_ = 0.002, *d*_3_ = 0.1046 and *d*_4_ = 0.0052, as defined by Hargrove (1994)[15].

We re-fit the model described by Lord et al., using the calibrated MODIS air temperature available for the area around Rekomitjie, to *G. pallidipes* catches between March 2000 and December 2016 using maximum likelihood estimation. We fitted the model to the data, fitting: i) only the parameters in the adult temperature-dependent mortality function (EQ3) (*a*_1_, *a*_2_); ii) only the parameters in the pupal temperature-dependent mortality function (EQ4) (*b*_1_, *b*_3_ and *b*_5_); and iii) fitting both functions. Parameter values were estimated using two iterations of the stochastic simulated annealing algorithm [32] followed by the Nelder-Mead algorithm [33]. We compared model fits for i-iii to the data using Akaike’s Information Criterion (AIC) [34]. The parameter values from the best fitting model were then used to implement separate closed-population ODE models for each 1 km x 1 km grid cell.

We used the fitted model to simulate tsetse population dynamics in each cell of the MODIS map using the adjusted MODIS air temperatures between March 2000 to December 2016. We did not know starting values for numbers of pupae and adults and therefore we used a ‘spin-up’ period of five years using temperature values from the first year in the series, allowing the model to stabilise before modelling populations from March 2000. We arbitrarily set the initial number of parous adults and pupae to 100, the number of nulliparous adults to 25, and the model was solved at monthly time steps. The simulation was performed as a closed-population model, with no movement through immigration or emigration of adjacent cells. We visualised predictions by combining estimates in each cell, at each time step, in a raster file [35]. The modelling process resulted in monthly spatial surfaces of abundance based on monthly spatial surfaces of adult mortality (*μ*_*A*_), pupal mortality (*μ*_*P*_), larviposition (*ρ*) and pupal emergence (*β*) rates for March 2000 – December 2016. We then generated mean surfaces for each metric for each year.

To compare modelled changes in population size over time, we produced two mean surfaces. One surface details the mean number of *G. pallidipes* per cell between 2001 and 2005; the second surface details the mean number of *G. pallidipes* per cell between 2012 and 2016. Averaging estimates over a 5-year period aimed to account for the effect of temperature trends such as the El Niño Southern Oscillation (ENSO) [36] and inter-annual variations on *G. pallidipes* populations.

## Results

### Comparison of MODIS and weather station temperatures

There was a mean difference of 2.1°C (SD = 1.264°C) between the land surface temperature from MODIS and air temperature from the Rekomitjie weather station. Conversion of MODIS LST to air temperature, using the Weiss calibration algorithm, reduced this difference to 0.831°C (SD = 0.611°C) at Rekomitjie. The linear model comparing Rekomitjie field station and converted MODIS data produced an *R*^2^ = 0.907, *RMSE* = 0.94. Comparisons across all 21 stations within Zimbabwe resulted in a mean difference of 1.094°C (SD = 0.823°C) (*R*^2^ = 0.901, *RMSE* = 1.56) (Fig 3) (data for individual stations is shown in Supplementary File 1).

**Figure 3:**
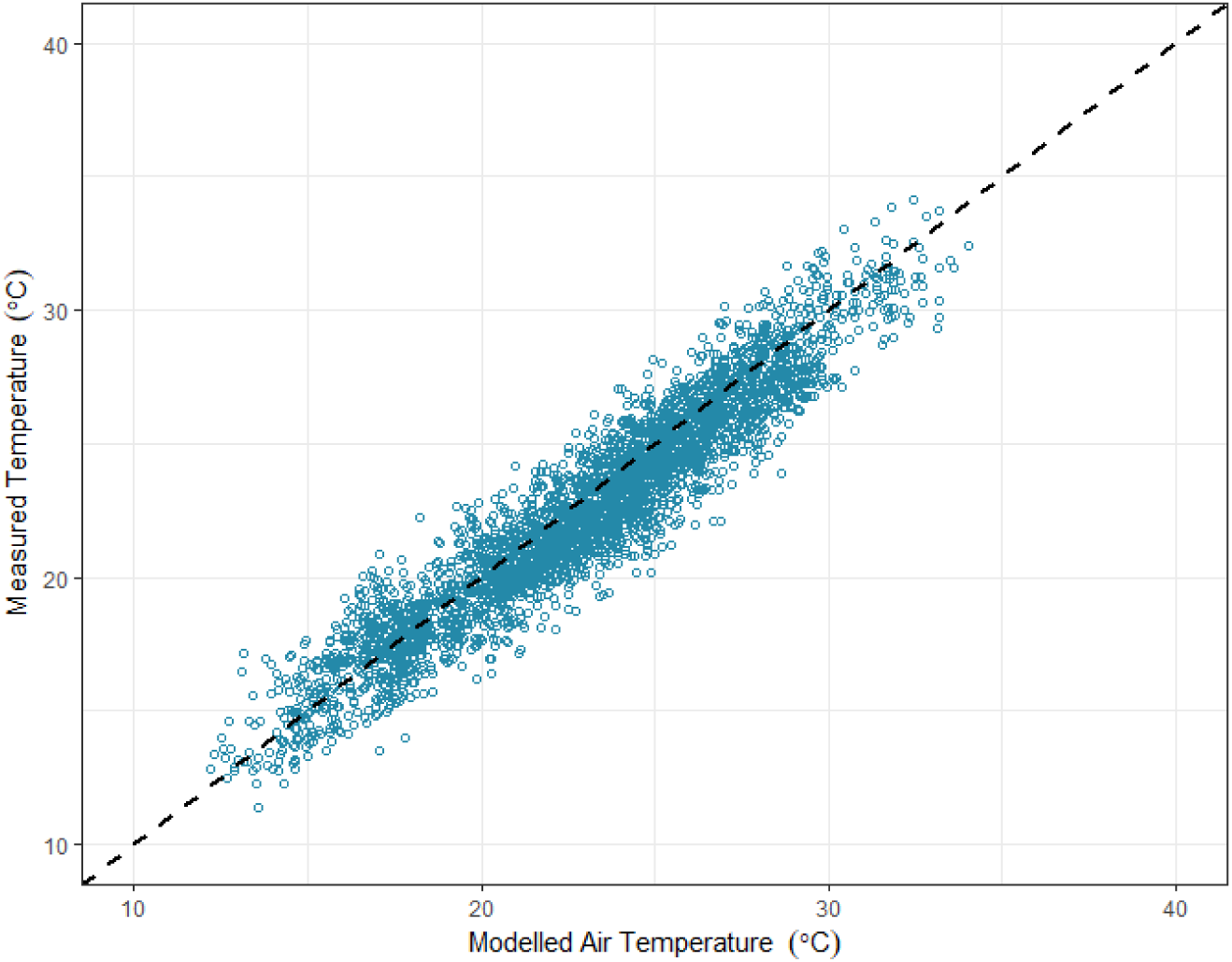
Comparison of monthly mean temperatures measured at weather stations and the same metric modelled from MODIS LST. Measured air temperatures were collected from 21 sites within northern Zimbabwe from 2000 to 2016. The dashed line represents the line of equality. *RMSE* = 0.94, *R*^2^ = 0.907.

### Population dynamics model

Optimisation of the initial parameter values for adult mortality (parameters *a*_1_ and *a*_2_), pupal mortality (parameters *b*_1_, *b*_3_ and *b*_5_) and density-dependent mortality (*δ*), produced a closer fit to the *G. pallidipes* catch data than the initial parameter values identified from the literature (AIC = 1746, Akaike weight (*w*(*AIC*)) = 4.83e^-68^). Allowing parameters in both the adult and pupal temperature-dependent mortality functions to vary, in addition to the density-dependent mortality coefficient, improved model fit (AIC = 1436, w(AIC) = 1), compared with only varying *δ* (AIC = 1746, w(AIC) = 4.83e^-68^), or only varying *δ* and either the adult mortality (AIC = 1465, w(AIC) = 3.11e^-07^) or the pupal mortality (AIC = 1789, w(AIC) = 2.22^e-77^) parameters. Final fixed and fitted parameter estimates for each function are shown in Supplementary File 2, alongside plots of the responses (Supplementary File 3). Our model using MODIS adjusted air temperatures was able to simulate the overall observed decline in *G. pallidipes* catches at Rekomitjie between 2000 and 2016 (Fig. 4) (*RMSE* = 4.78, *R*^2^ = 0.65).

**Figure 4:**
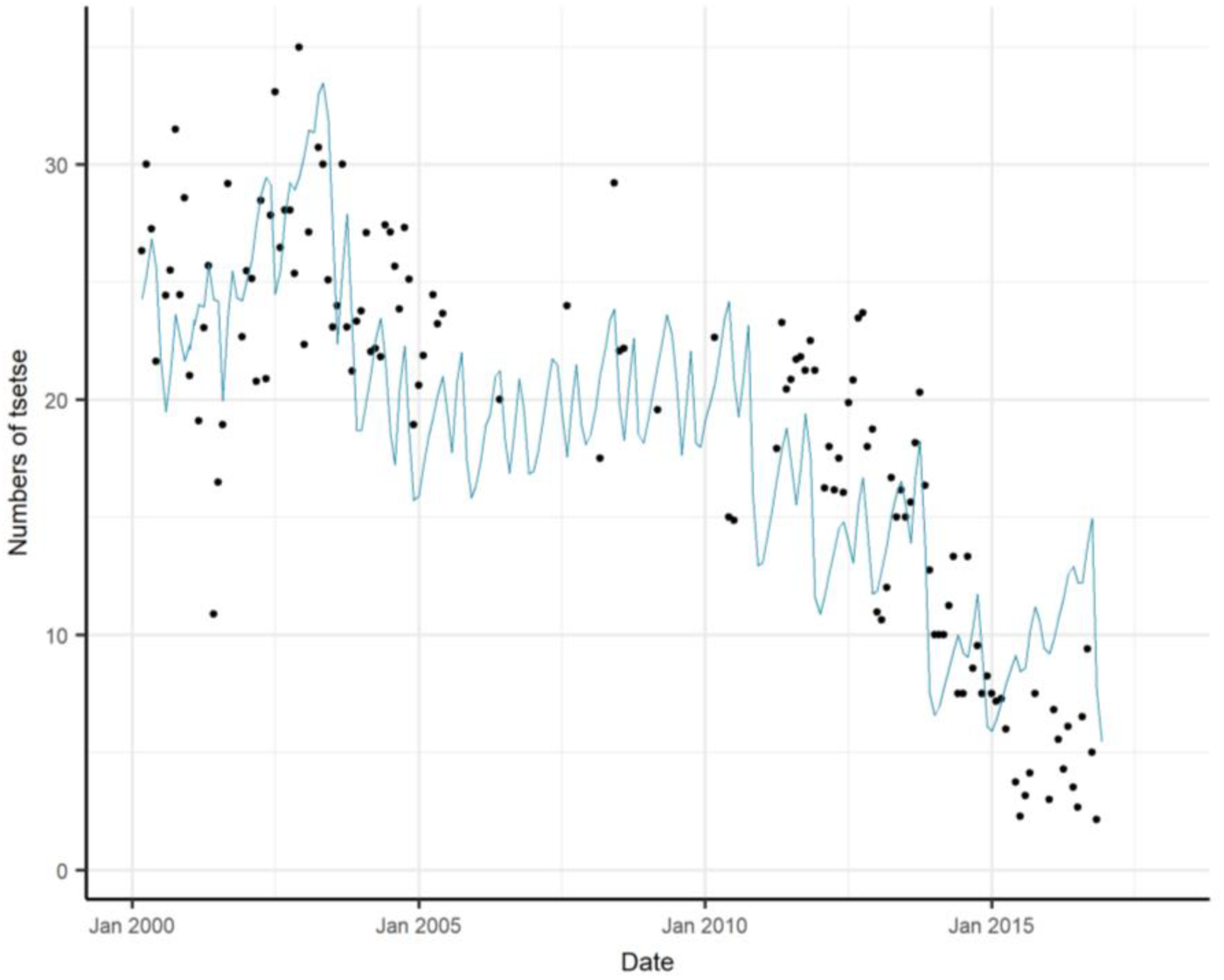
Observed (black dots) and modelled (blue line) changes in number of *G. pallidipes* females caught at Rekomitjie between March 2000 and December 2016. The model described by Lord et al. [19] was refit to tsetse catch data utilising MODIS adjusted air temperatures. Black dots represent the average number of tsetse caught per month at Rekomitjie, the blue line represents model fit.

### Spatial projection of population dynamic model

Combining the closed population models resulted in surfaces detailing spatial variation in daily adult mortality, daily pupal mortality, larviposition rate and daily pupal emergence rate across Northern Zimbabwe. When comparing estimates of mean adult population size across years (Fig. 5A and 5B), several high population density areas occurring within 2001-2005 estimates are predicted to have decreased in 2012-2016, for example locations within the north of Mashonaland Central Province and Mashonaland West Province, within the Zambezi valley.

**Figure 5.**
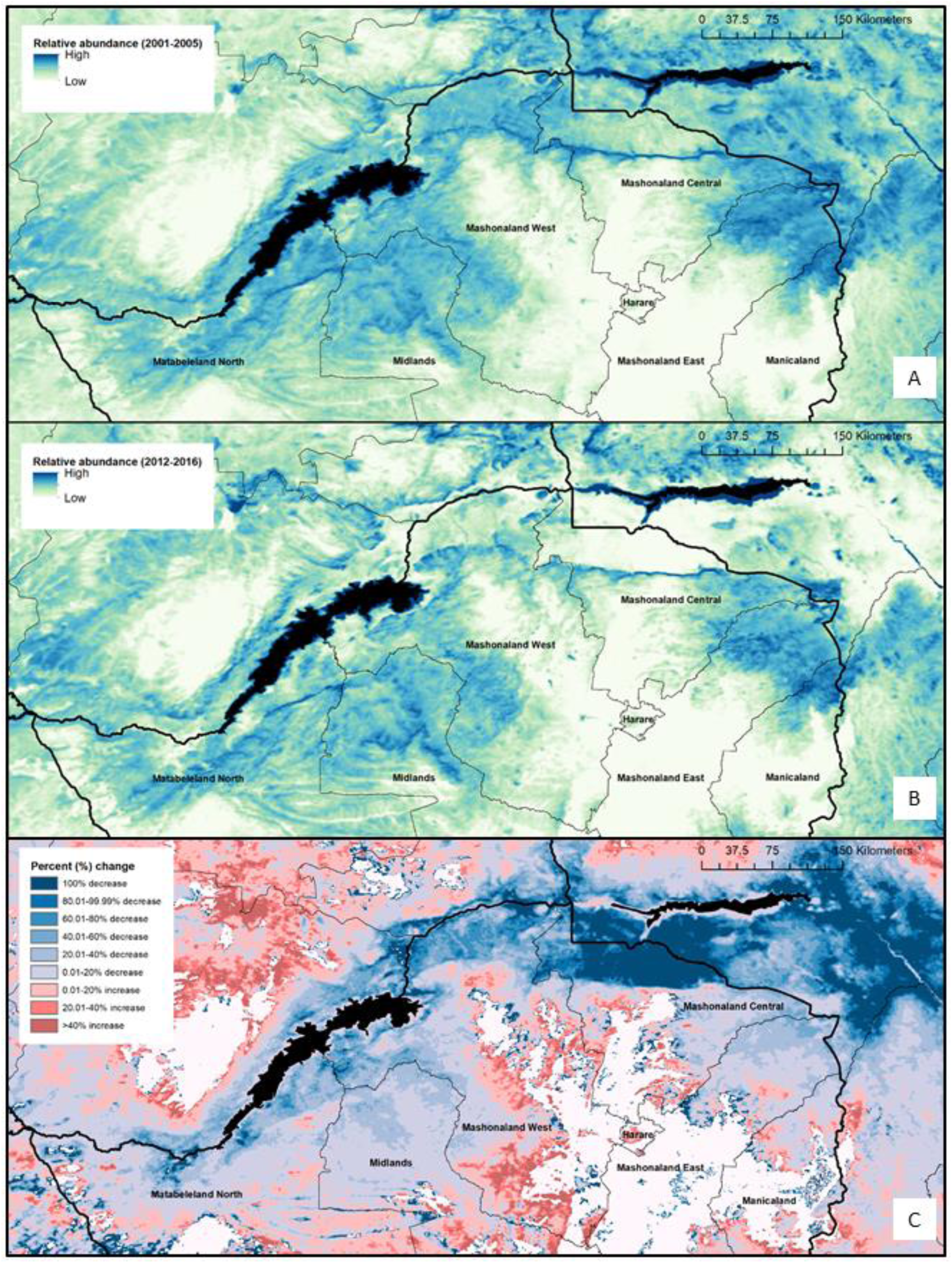
Spatial variation within the estimated relative abundance of *G. pallidipes*. A) Relative abundance of *G. pallidipes* within Northern Zimbabwe (mean across 2001-2005); B) Relative abundance of *G. pallidipes* within Northern Zimbabwe (mean across 2012-2016); C) Relative difference between estimated abundance (2001-2005 vs 2012-2016). Dark blue indicates areas of decrease in predicted abundance, whereas red indicates areas of a predicted increase in abundance. Areas of no change (either stable populations, or non-suitable environments) are shown in white.

Generally, the spatial predictions suggest an overall decrease in *G. pallidipes* population density over time, except for a few locations within Midlands Province, and Matabeleland North Province, which are predicted to have become more suitable for tsetse in terms of temperature. The overall pattern, when compared with an elevation surface (Fig. 1), appears to indicate a shift in tsetse populations from lower elevation areas to higher elevation areas, indicating increased suitability at higher altitudes (∼1000 M.A.S.L, Fig. 6). A categorical surface showing percentage change in relative abundance between the two periods is provided as Fig. 5C to aid interpretation of the modelled population density surfaces shown. Fig. 5C helps to identify several areas in which tsetse populations may have remained stable over time; these areas are primarily areas in which populations have remained unsuitable in terms of temperature – for example several areas in Mashonaland East and Manicaland Provinces, alongside a hotspot within Matabeleland North province.

**Figure 6.**
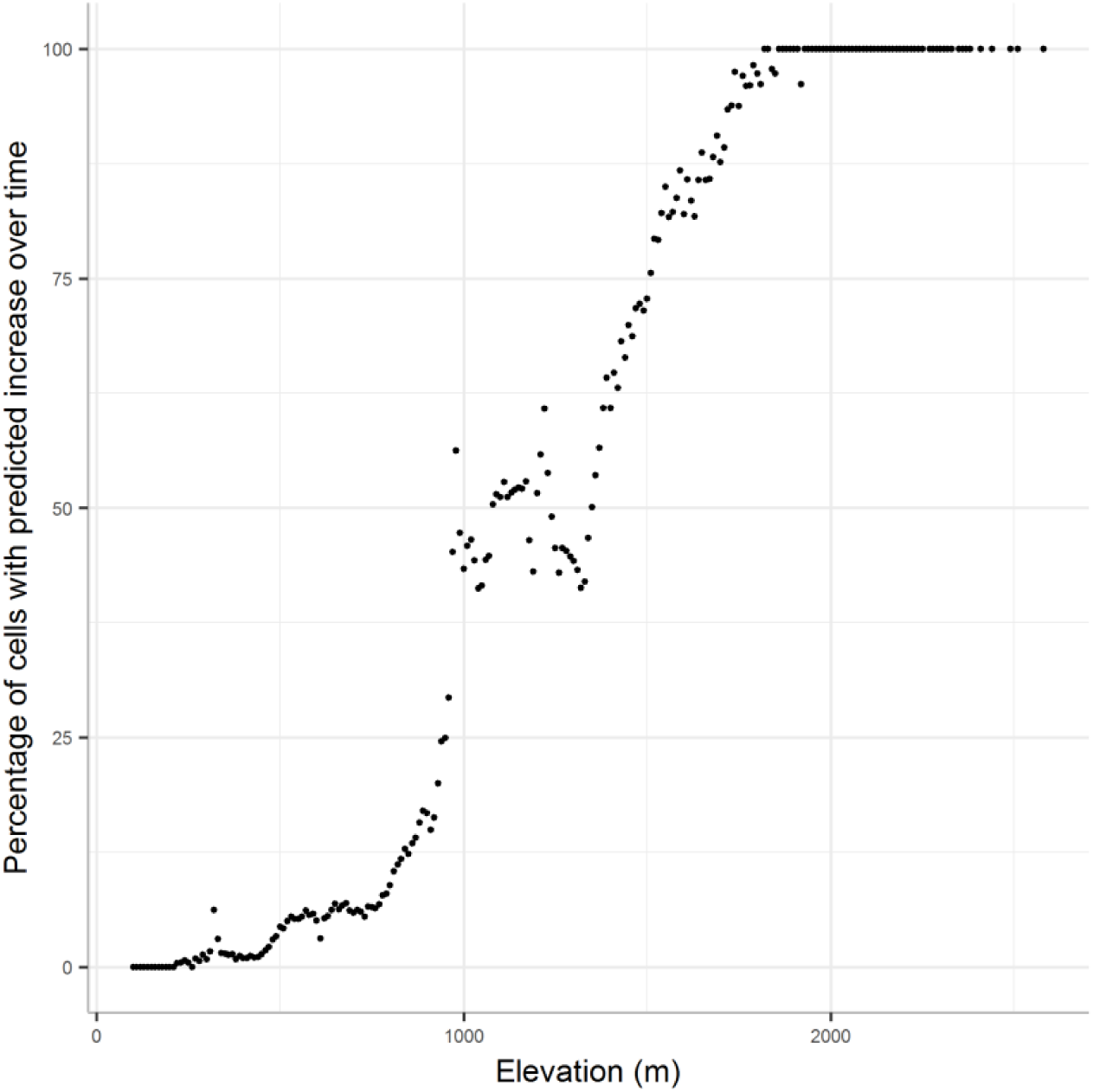
Percentage of cells showing a predicted increase in number of tsetse when comparing mean 2001-2005 with mean 2012-2016, per elevation class.

When investigating trends based by elevation (Fig. 6), it appears that most locations for which relative abundance is predicted to increase occur at high elevation locations ∼1000 M.A.S.L, with >50% of locations at this altitude predicted to have increasing population counts across the study period.

## Discussion

Understanding the distribution and abundance of tsetse flies is crucial for the identification of remaining HAT and AAT foci, and subsequent targeting of control strategies in the field. Here, we build on our original finding, that temperature increases explain a >90% decline in the *G. pallidipes* population at one site in Zimbabwe, to show that while a general decline in tsetse populations across northern Zimbabwe may have occurred, there may be areas at higher elevation (>1000 M.A.S.L) where temperature has become more suitable.

Although the kind of data used to parameterise this model are relatively rare, our work highlights that recent climate change is probably having an impact on tsetse population dynamics in sub-Saharan Africa and may already be changing the distribution of disease, particularly for AAT [37]. We show here, as a proof of concept, how longitudinal catch data can be combined with other metrics to quantify r-HAT vector abundance; advocating for the collection, collation, and sharing of data on tsetse abundance at other HAT endemic sites within sub-Saharan Africa.

Our MODIS adjusted air temperature surfaces showed close alignment with temperatures recorded at weather stations across northern Zimbabwe, providing confidence in the use of these data to inform models for locations where meteorological data was lacking. The initial model adapted from Lord *et al*. provided a good fit to the field obtained count data for *G. pallidipes* at Rekomitjie.

Unfortunately, the lack of access to longitudinal catch data for other locations within the study extent meant that methods of model validation were limited. Future work should focus on model validation, via collation and utilisation of additional longitudinal catch data, should they exist, or in-field sampling in locations predicted to be areas of high and low abundance.

The use of monthly mean temperature data within this analysis potentially mask daily fluctuations in temperature, as each life stage is exposed to a constant temperature for the duration of each time step (30 days). Although there is uncertainty surrounding *in situ* temperature exposure of tsetse, future work could investigate the use of 8-daily imagery and interpolation methods to reduce the time step in the modelling process from 30 to 8 days [22]. This finer temporal resolution may better represent *in situ* temperature fluctuations, however, poses further computational constraints, and micro-temporal fluctuations may remain uncharacterised. Diurnal temperature cycles (*Air*_*max*_ – *Air*_*min*_) have been shown to have a large impact on other disease vectors, for instance several *Anopheline* species [38].

The model we used (Lord *et al*. [19]) is constructed using parameters for female fly survival, larviposition rates, and pupal emergence rates. Previous work has identified that male flies may experience periods of sterility when exposed to sustained temperatures exceeding 30°C [39], furthermore, variation in pupal development rates occur between sexes, with female flies emerging 1-2 days before males in laboratory colonies [29]. There may be additional processes affected by temperature that are not captured within our modelling framework, and the inclusion of these additional parameters may prove non-trivial.

Another thing to bear in mind is that the training data, and consequently the predictions of abundance, relate to a tsetse population under no exposure to control. Further information is required regarding current vector control status within cells outside of Rekomitjie, as the presence of control will dramatically influence tsetse abundance [40]. Control operations pre-2000 are well documented and widespread across Zimbabwe [3, 41, 42], however, unfortunately there is little published literature detailing interventions applied post-2000. In historic instances, following from tsetse control operations in or near the Zambezi valley (for example Muzarabani and Dande), there were settlement and land-use changes. These changes would potentially result in a reduced ability of tsetse to recover in these areas if both the habitat has degraded and temperature has increased. An interesting application of the model produced here would be to incorporate ‘control events’ at specific time periods, by the introduction of an additional control parameter, to investigate the local scale effects of the introduction/or scale up of vector control interventions. Model manipulation would allow for spatial predictions of intervention efficacy, highlighting locations where specific vector control techniques can be employed to exploit spatial sensitivities in tsetse dynamics.

This work should be interpreted in the context of several key limitations. To generate the spatial predictions described here, each 1 km × 1 km cell was processed as a separate closed population with no immigration or emigration occurring across cells. There are several known limitations to this approach, with, for some populations, immigration and emigration being more determinant than births and deaths. With the absence of dispersal in this model, an extinction event captured by the ODE renders a cell inhospitable for future occupancy. There is a general agreement that periodic local extinctions and recolonisations are common in nature [43], therefore future work should investigate the construction of a metapopulation dynamic model, in which local populations can interact via dispersal events. Quantifying tsetse interaction and movement would aid modelling the effects of interventions discussed above, with current estimates of tsetse dispersal being in the region of ∼350m per day [44].

Our analyses suggest a shift in tsetse populations from areas of lower elevation, to areas of higher elevation. Such predictions support theories that certain areas within the Zambezi valley will soon be too hot to support populations of *G. pallidipes* [19], and areas previously considered too cold for tsetse would become more environmentally suitable if climate trends continue. Our analysis, however, only considers temperature. There may have been other environmental changes related to settlement and land-use which may also contribute to a decline in tsetse numbers within our study extent [45]. Further work is required to investigate the effect of these factors. Additionally, areas predicted to have become more suitable for tsetse may lack suitable habitat or host densities to support viable tsetse populations. Interestingly, most recent cases of r-HAT in Northern Zimbabwe have come from the vicinity of Makuti, which is in an area where our model predicts an increase in climatic suitability for tsetse [27], suggesting a suitable environment for parasite and hosts in this area.

Lastly, tsetse presence alone is not indicative of HAT or AAT risk; for transmission of sleeping sickness to occur, there is a need for the triad of parasite, vectors and hosts [46, 47]. Human behaviour at various scales also contributes to disease transmission and the effects of climate will impact not only on tsetse but also hosts and human activities. While we highlight how climate change may influence tsetse distribution and abundance, there will be complex interactions which may exacerbate or mitigate disease risk. Ultimately, quantifying the presence and prevalence of these other factors will allow for estimates of remaining disease foci within Northern Zimbabwe, resulting in a more refined public health tool.

## Ethics approval and consent to participate

Not applicable.

## Consent for publication

Not applicable.

## Availability of data and material

Model code can be accessed at https://github.com/jenniesuz/tsetse_climate_change.

## Authors’ contributions

Conceptualization: Stephen J. Torr, Joshua Longbottom, Jennifer S. Lord.

Data curation: Stephen J. Torr, Jennifer S. Lord, Joshua Longbottom, Harry S. Gibson, Daniel J. Weiss. Formal analysis: Joshua Longbottom.

Funding acquisition: Joshua Longbottom, Stephen J. Torr. Methodology: Joshua Longbottom, Jennifer S. Lord.

Supervision: Stephen J. Torr, Jennifer S. Lord. Writing – original draft: Joshua Longbottom.

Writing – review & editing: Joshua Longbottom, Stephen J. Torr, Jennifer S. Lord, Cyril Caminade, Harry S. Gibson, Daniel J. Weiss.

## Acknowledgements

We wish to express our sincerest gratitude for the data produced and provided by the Tsetse Control Division of the Zimbabwe Public Service. We thank Dr Glyn Vale for his extremely valuable comments on the manuscript.

