## Supplementary figures and images for "Modelling the impact of climate change on the distribution and abundance of tsetse in Northern Zimbabwe"

### Supplementary File 1

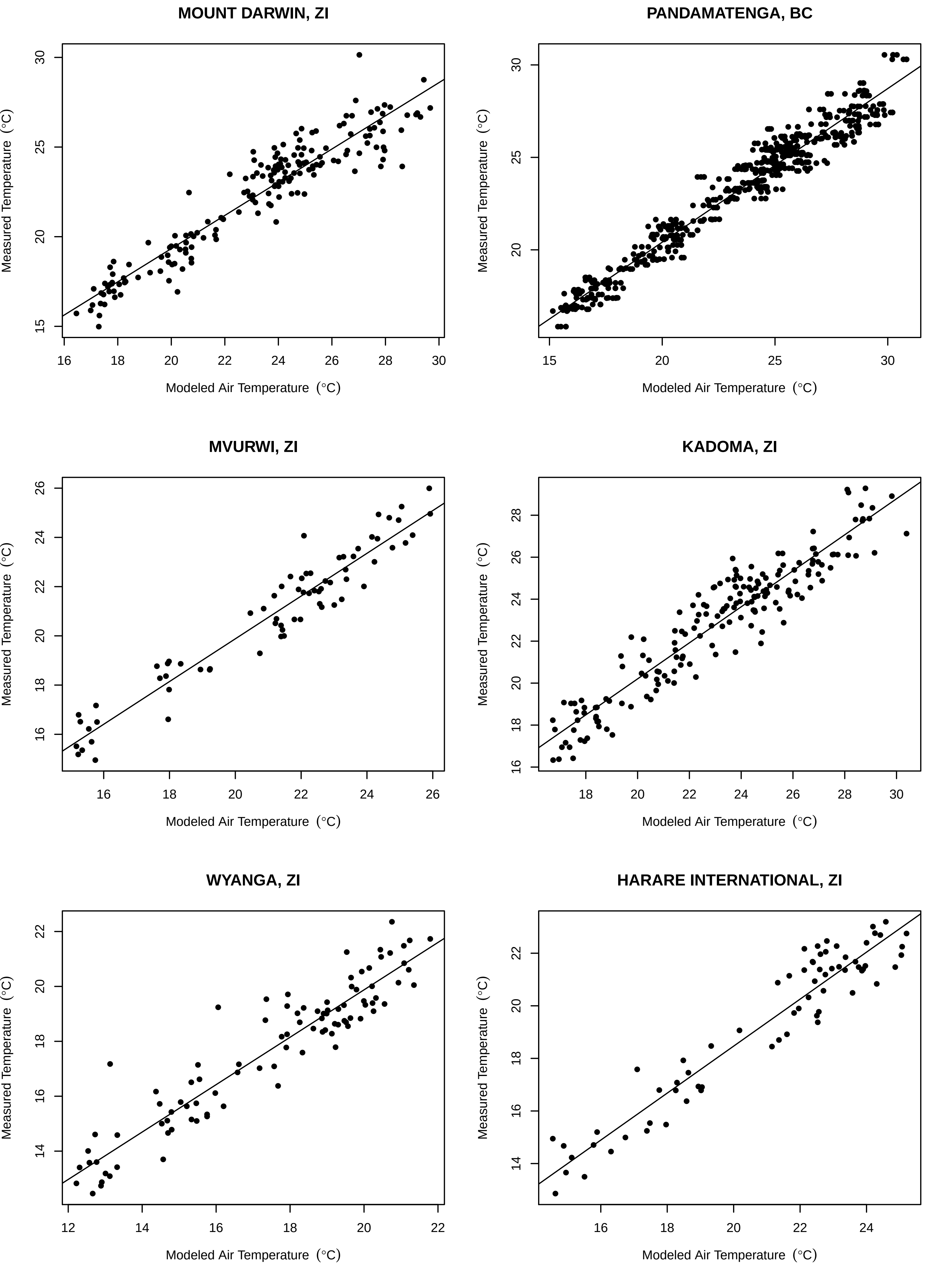

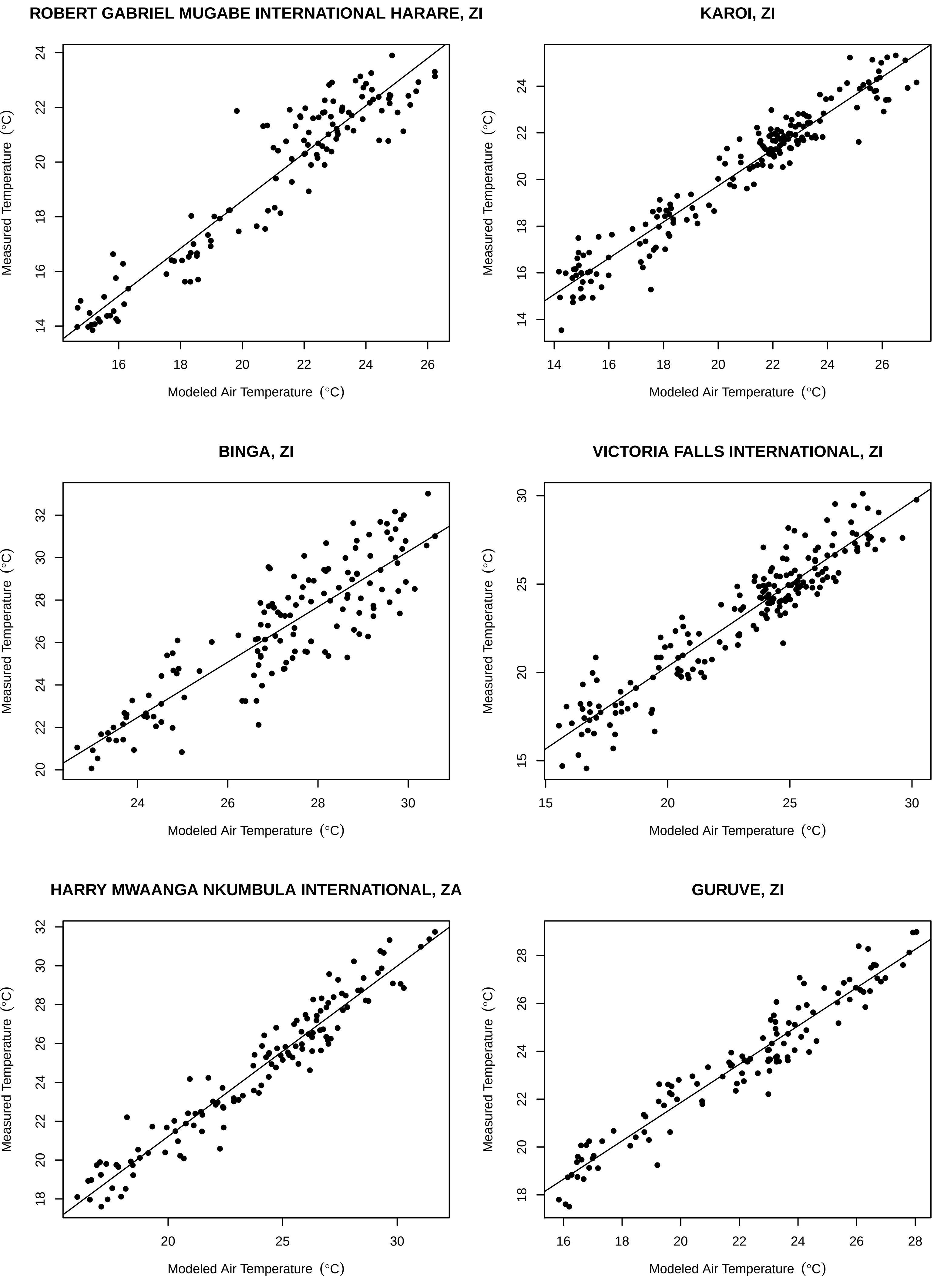

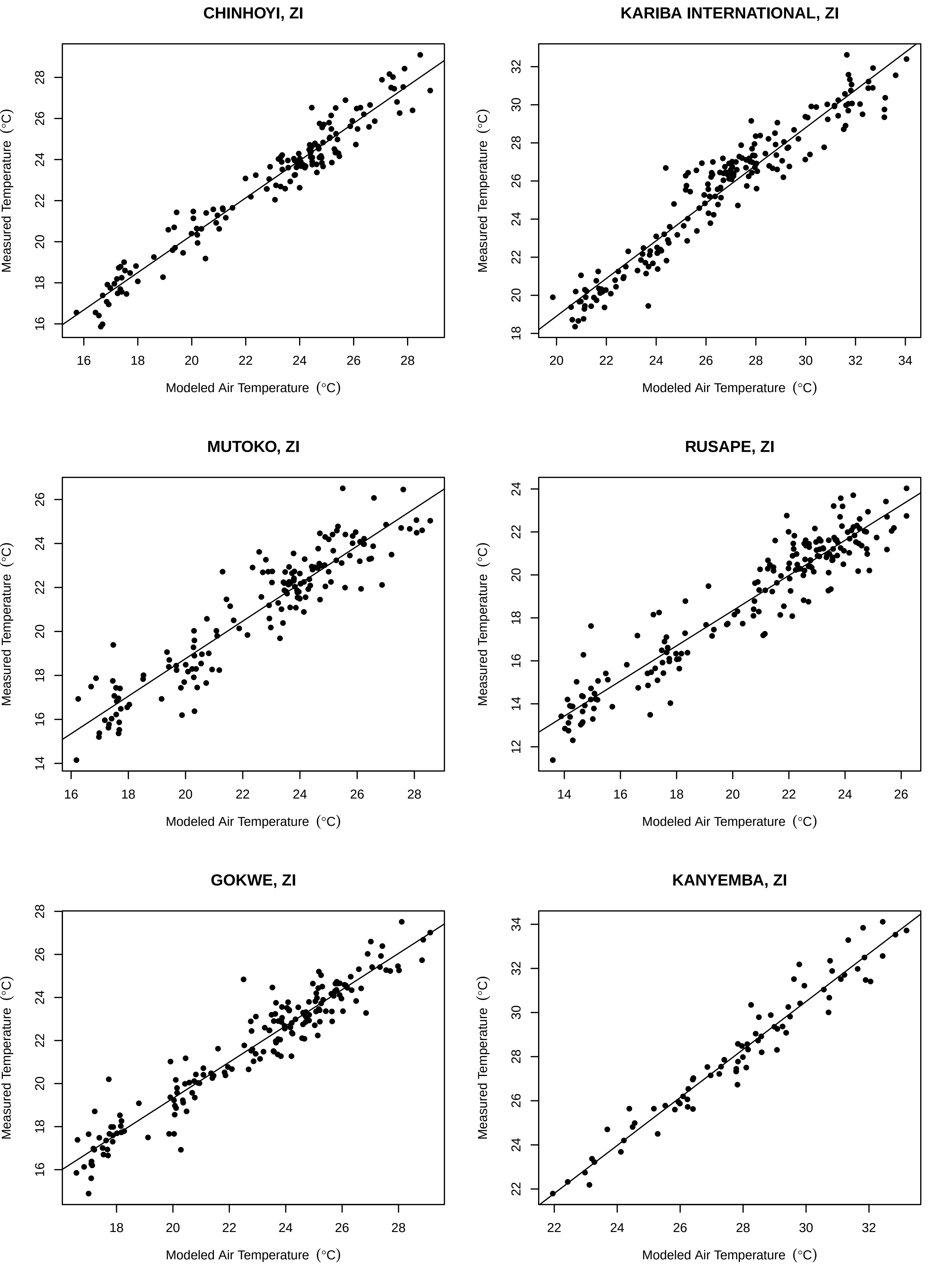

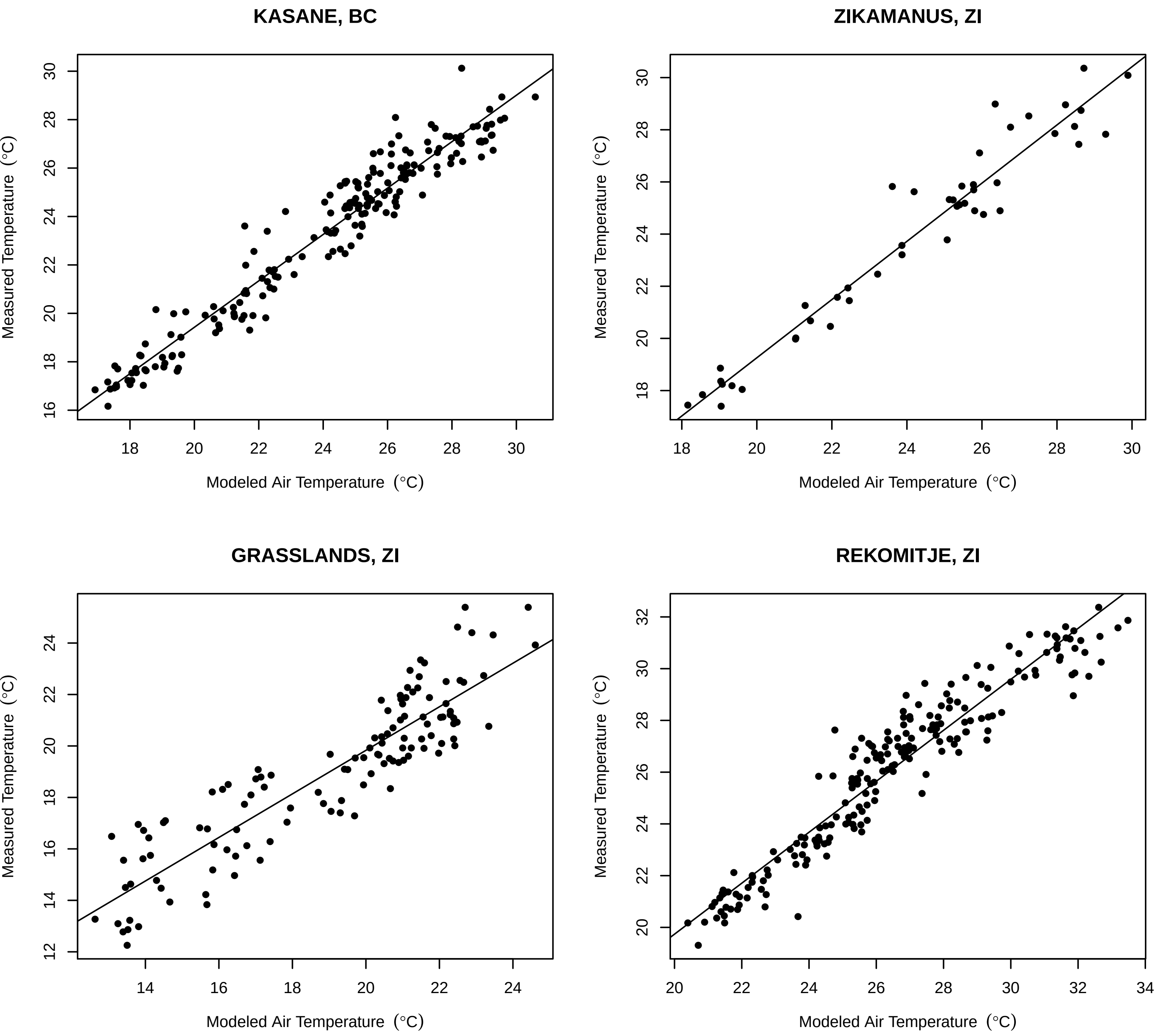
