## Supplementary File 2 for "Modelling the impact of climate change on the distribution and abundance of tsetse in Northern Zimbabwe"

**Summary of model parameters.** Table contains both fixed and estimated parameter values from the population dynamic model with the lowest AIC.

| **Parameter** | **Function or parameter definition** | **Estimate from fit of Eq. 1-4 to published laboratory and field data** | **Estimate from fit of population dynamic model** |
| --- | --- | --- | --- |
| *a_1_* | Eq. 3: Adult mortality rate (*µ_A_*) as a function of temperature | 0.027 ± 0.001 | 0.0369 ± 3.91e^-5^ |
| *a_2_* |  | 0.153 ± 0.020 | 0.0452 ± 1.51e^-4^ |
| *b_1_* | Eq. 4: Pupal mortality rate (*µ_P_*) as a function of temperature | 0.0019 ± 0.0004 | 0.00035 ± 1.98e^-6^ |
| *b_2_* |  | 0.006 ± 0.001 | Fixed |
| *b_3_* |  | 1.481 ± 0.681 | 1.1183 |
| *b_4_* |  | 0.003 ± 0.001 | Fixed |
| *b_5_* |  | 1.211 ± 0.117 | 0.8056 ± 3.04e^-3^ |
| *c_1_* | Eq. 5: Pupal emergence rate ($\beta$) | 0.05884 ± 0.00289 [2] | Fixed |
| *c_2_* |  | 4.8829 ± 0.0993 [2] | Fixed |
| *c_3_* |  | -0.2159 ± 0.0050 [2] | Fixed |
| *d_1_* | Eq. 6: Larviposition rate (*ρ*_n_) | Nulliparous - 0.061 ± 0.002 [1] | Fixed |
| *d_2_* |  | Nulliparous - 0.002 ± 0.0009 [1] | Fixed |
| *d_3_* | Eq. 7: Larviposition rate (*ρ_p_*) | Parous - 0.1046 ± 0.0004 [1] | Fixed |
| *d_4_* |  | Parous - 0.0052 ± 0.0001 [1] | Fixed |
| *δ* | Density-dependent mortality coefficient | NA | 0.000105 ± 3.56e^-7^ |

1.	Hargrove JW. Reproductive rates of tsetse flies in the field in Zimbabwe. Physiological Entomology. 1994;19 4:307-18; doi: doi:10.1111/j.1365-3032.1994.tb01057.x. <https://onlinelibrary.wiley.com/doi/abs/10.1111/j.1365-3032.1994.tb01057.x>.

2.	Phelps RJ, Burrows PM. Puparial duration in *Glossina mortisans orientalis* under conditions of constant temperature. Entomologia Experimentalis et Applicata. 1969;12 1:33-43; doi: doi:10.1111/j.1570-7458.1969.tb02494.x. <https://onlinelibrary.wiley.com/doi/abs/10.1111/j.1570-7458.1969.tb02494.x>.
