## Supplementary File 3 for "Modelling the impact of climate change on the distribution and abundance of tsetse in Northern Zimbabwe"

**Fitted temperature-dependent functions.**

(a) Adult female mortality rate per day: points—published estimates from mark-recapture experiments on Antelope Island, Zimbabwe [1]; line—fitted temperature-dependent adult mortality function (Eq 3). (b) Pupal mortality rate per day: points—published estimates from laboratory experiments [2]; line—fitted temperature-dependent pupal mortality function (Eq 4). (c) Pupal emergence rate per day: points—published estimates from laboratory experiments; line—Eq 5 fitted as described in [3]. (d) Larviposition rate per day: points—data from published field experiments [4]; lines—Eq 7 fitted as described in [1].

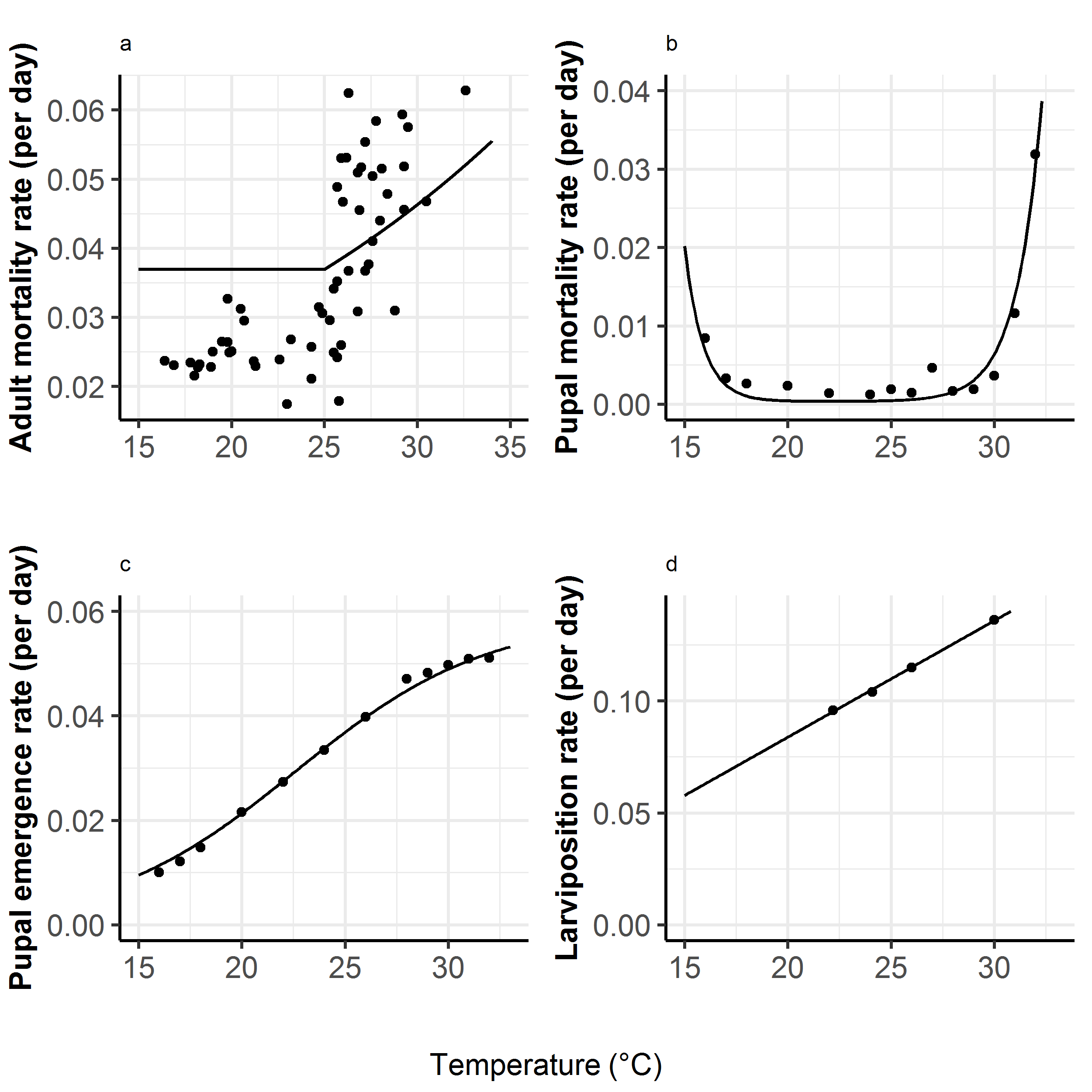

1.	Hargrove J. Tsetse population dynamics. In: Maudlin I, Holmes PH, Miles MA, editors. The trypanosomiases; 2004.

2.	Phelps RJ. The effect of temperature on fat consumption during the puparial stages of *Glossina morsitans morsitans* Westw. (Dipt., Glossinidae) under laboratory conditions, and its implication in the field. Bulletin of Entomological Research. 1973;62 3:423-38; doi: 10.1017/S0007485300003953. <https://www.cambridge.org/core/article/effect-of-temperature-on-fat-consumption-during-the-puparial-stages-of-glossina-morsitans-morsitans-westw-dipt-glossinidae-under-laboratory-conditions-and-its-implication-in-the-field/EBA7AB997B436C038C5FDB190750B601>.

3.	Phelps RJ, Burrows PM. Puparial duration in *Glossina mortisans orientalis* under conditions of constant temperature. Entomologia Experimentalis et Applicata. 1969;12 1:33-43; doi: doi:10.1111/j.1570-7458.1969.tb02494.x. <https://onlinelibrary.wiley.com/doi/abs/10.1111/j.1570-7458.1969.tb02494.x>.

4.	Hargrove JW. Reproductive rates of tsetse flies in the field in Zimbabwe. Physiological Entomology. 1994;19 4:307-18; doi: doi:10.1111/j.1365-3032.1994.tb01057.x. <https://onlinelibrary.wiley.com/doi/abs/10.1111/j.1365-3032.1994.tb01057.x>.
